# Identification of polyphosphate-binding proteins in *E. coli* uncovers targets involved in translation control and ribosome biogenesis

**DOI:** 10.1101/2025.02.12.637445

**Authors:** Kanchi Baijal, Brianna Kore, Iryna Abramchuk, Alix Denoncourt, Shauna Han, Abby Simms, Amy Dagenais, Abagail R. Long, Adam D. Rudner, Mathieu Lavallée-Adam, Michael J. Gray, Michael Downey

## Abstract

In many bacteria, polyphosphate kinase (PPK) enzymes use ATP to synthesize polyphosphate (polyP) in response to cellular stress. These chains of inorganic phosphates are joined by high-energy bonds and can reach hundreds of residues in length. PolyP plays diverse functions in helping bacteria adjust to changing environmental conditions. However, the molecular mechanisms underlying these functions are poorly understood. In eukaryotic cells, polyacidic serine- and lysine-rich (PASK) motifs of proteins can mediate binding to polyP chains. Whereas PASK motifs are relatively common in yeast and human cells, we report that these sequences are rare in bacteria commonly used for polyP research. Thus, to identify novel polyP-binding proteins in *Escherichia coli,* we carried out an untargeted screen and identified 7 novel targets with links to translation control and ribosome biogenesis. For two targets, the GTPase activating protein YihI and the ribonuclease Rnr, we mapped the regions of polyP interaction to non-PASK sequences and identified lysine residues critical for binding. We found that deletion of *rnr* suppressed the slow growth phenotype of Δ*ppk* mutants grown on minimal media. Conversely, *ppk* deletion resulted in decreased Rnr protein expression. These phenotypes were dependent on the polyP binding region of Rnr but independent of polyP binding itself, suggesting a complex interplay between PPK and Rnr function in *E. coli.* Overall, our work provides new insights into the scope of polyP binding proteins and extends the connections between polyP and the regulation of protein translation in *E. coli*.

## INTRODUCTION

Polyphosphate (polyP) chains are multifunctional polymers composed of three to hundreds of phosphate monomers linked by high-energy phosphoanhydride bonds^1^. Although polyP molecules are found broadly across prokaryotic and eukaryotic cells, the mechanisms of polyP synthesis differ between species^2^. In bacteria, polyP is synthesized by the polyphosphate kinase (PPK) enzymes, usually in response to cellular stresses such as starvation^3^ or treatment with oxidizing agents^4^. While some bacteria, such as the Gram negative *Escherichia coli,* have only one PPK enzyme, others express both PPK1 and PPK2 proteins^2^. Compared to PPK2, PPK1 is the dominant polyP-synthesizing enzyme in bacteria and preferentially uses ATP as a substrate^5^. PPK1 enzymes can also catalyze the reverse reaction to generate ATP from ADP and polyP^6^. although the extent to which this activity regulates pools of polyP *in vivo* is uncertain. Alternatively, polyP molecules can be degraded into free inorganic phosphate (Pi) via the action of the exopolyphosphatase PPX, which cleaves phosphoanhydride bonds beginning at the ends of polyP chains^7^. Bacterial cells mutated for *ppk* genes show defects in stress and antibiotic resistance^4, 8, 9^, reduced biofilm formation^10^, and decreased ability to infect host cells^11^. These phenotypes underly efforts to develop PPK inhibitors as a new tool in the fight against antimicrobial resistance. PPK enzymes are also present in some lower eukaryotic organisms, including the slime mold *Dictyostelium discoideum*, having been acquired by horizontal gene transfer^12, 13^.

In yeast, polyP is synthesized by the vacuole-bound vacuolar transporter chaperone (VTC) complex^14^. VTC activity is coupled to polyP transport into the vacuole lumen and its sequestration therein^15^. The VTC complex (and presumably polyP) has been linked to ion and phosphate homeostasis^16, 17^, cell cycle control^18^, microautophagy^19^, and the regulation of protein translation^20^. There are no mammalian homologs of VTC or PPK proteins, and the mechanism of polyP synthesis remain poorly defined in higher eukaryotes such as humans^21, 22^. There is one report that the mitochondrial F_o_F1 ATPase can synthesize polyP^23^, but it is unclear if this activity impacts total cellular levels of the polymer. The levels of polyP in human cells are generally thought to be lower than that measured in microorganisms^21^, although this assertion has recently been challenged^24^. Regardless, diverse roles for polyP have been suggested in mammalian cells including cell signaling^25–27^, protein folding^28^, energy metabolism^29^, and blood clotting^30^. While polyP could impact cell function through diverse mechanisms, there is particular interest in roles mediated by its interaction with protein targets (reviewed in^31^). Previous work in eukaryotes has collectively identified dozens of polyP-binding partners^20, 32–37^. In bacteria, however, examples of polyP-interacting proteins are less common. In *E. coli*, during stress, polyP serves as a molecular adaptor for the Lon protease to promote the degradation of ribosomal proteins as well as the DnaA replication initiation protein^38, 39^. PolyP binding to CsgA plays a role in the regulation of biofilm formation^28^. Finally, polyP also binds to the chaperone Hfq to promote its tight interaction with DNA and regulate its phase separation^40^. Beyond *E. coli*, the regulation of stress responses by polyP binding proteins is a common theme. For example, in *Helicobacter pylori*, polyP binding to sigma 80 is thought to directly regulate a transcriptional program to help bacteria adapt to starvation^41^. Since the deletion of *ppk* homologs in many bacteria impacts diverse molecular pathways^10, 40, 42, 43^, we speculated that additional polyP binding proteins remain to be found.

In this work, we report the use of an untargeted proteomic screen to identify 7 novel polyP-binding proteins in *E. coli.* Remarkably, all 7 of these targets are linked to ribosome biogenesis and protein translation. For two proteins, YihI and Rnr (RNase R), we mapped the region of polyP binding to lysine-rich sequences of the proteins that are important for ribosome- and translation-related functions. Unexpectedly, while Rnr levels are downregulated in Δ*ppk* mutants relative to wild-type controls grown on minimal media, deletion of the *rnr* gene or truncation of the polyP-binding region suppresses the slow growth phenotype of Δ*ppk* mutants under these same conditions. However, mutational analysis revealed that these effects are likely independent of Rnr binding to polyP, suggesting the possibility that polyP impacts Rnr function through both direct and indirect mechanisms. Together, our work extends the scope of polyP-protein interactions in *E. coli* and identifies new avenues for exploration of the PPK-dependent regulation of ribosome biogenesis and protein translation *in vivo*.

## RESULTS

### The landscape of PASK-containing proteins in bacteria

In eukaryotic cells, we have been particularly interested in the interaction of polyP with polyacidic-serine and lysine-rich (PASK) motifs of target proteins. In *Saccharomyces cerevisiae*, for example, there are 427 PASK-containing proteins, and work from our group and others have validated polyP binding to 27 of these^20, 36, 44^. While interaction between polyP and PASK-containing proteins was originally proposed to be covalent^36^, recent work challenges this assertion, suggesting instead a non-covalent interaction with positively charged PASK lysines^34^. Regardless of the mechanism at play, we reasoned that PASK-containing proteins would be excellent candidates for novel polyP effectors in bacteria.

To investigate the occurrence of PASK motifs in bacteria, we searched the proteomes of both Gram negative and positive species commonly used in polyP research. We did so using a program we call PASKMotifFinder, which was also recently used to find PASK-containing proteins in Trypanosomes^45^. We defined a PASK motif as a protein subsequence of 20 amino acids containing at least 75% D/E/S/K residues and at least one lysine residue, consistent with the definition we used previously for eukaryotic cells^20^. We found that compared to yeast and human cells, PASK-containing proteins are rare in both reviewed (**Figure 1A**) and unreviewed (**Figure S1A**) UniProt database entries^46^ from proteomes of bacterial species commonly used for polyP research.

**Figure 1.**
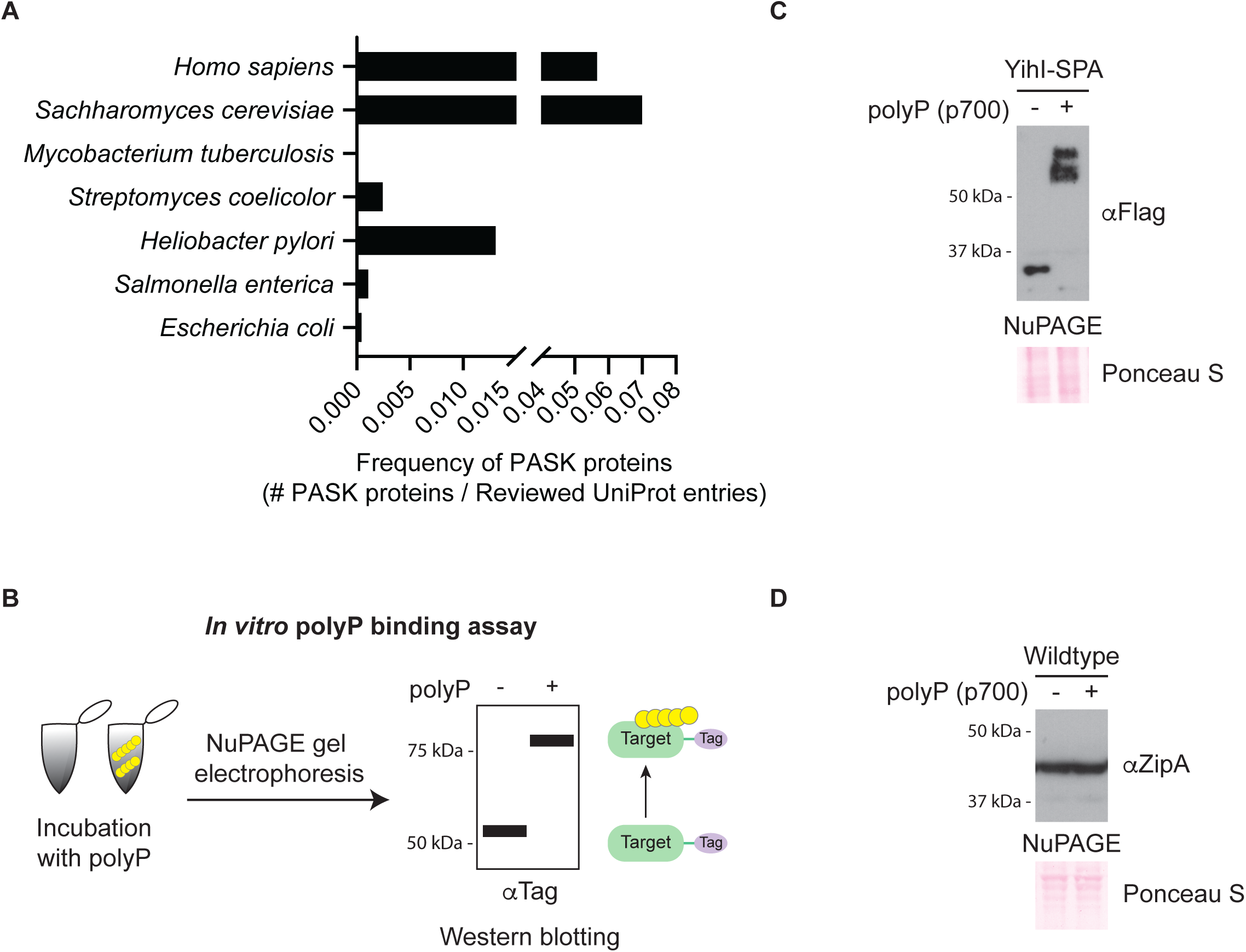
Characterization of PASK sequences in *E. coli.* **(A)** Frequency of PASK motifs in bacteria. The number of proteins containing 1 or more PASK motifs (75% D/E/S/K content with at least one lysine within a 20 amino acid window) from reviewed proteomes of the indicated species were normalized by the total number of reviewed UniProt entries of each species. **(B)** Schematic of the *in vitro* polyP binding assay. Whole cell extracts incubated in the absence or presence of synthetic polyP (p700) were resolved using a Bis-Tris gel (sold under the name NuPAGE) electrophoresis. Target proteins were visualized by western blotting using an antibody towards an epitope tag or the endogenous protein. Proteins that have slower migration in the presence of polyP compared to in its absence are thought to bind polyP. **(C-D)** *In vitro* polyP binding to (C) YihI-SPA and (D) ZipA. Assays were conducted as described in B. In both cases, samples were resolved using NuPAGE and transferred to PVDF. YihI-SPA and ZipA were detected using anti-Flag or anti-ZipA antibodies, respectively. Ponceau S was used to show that samples migrated equally. Images are representative of results from ≥3 experiments.

### *E. coli* YihI is a novel polyP binding protein

We focused on the only two PASK-containing proteins in *E. coli*, ZipA and YihI, that were identified using PASKMotifFinder. ZipA is an essential protein required for cell division^47^, and YihI is an activating protein for the essential GTPase Der^48^. Together, these two proteins represent 0.05% of the total proteome – a stark contrast to the situation in *S. cerevisiae*, where the fraction of PASK-containing proteins is 7.3%^20^. To determine if ZipA and YihI interact with polyP, we looked for polyP-induced electrophoretic shifts (hereafter ‘polyP shift’) on bis-tris polyacrylamide gels, which are sold commercially under the NuPAGE brand name (**Figure 1B**). This technique has previously been used to characterize polyP binding to PASK motifs^20, 34, 36, 49^. To conduct these *in vitro* polyP binding assays, we incubated whole cell extracts from SPA-tagged and wild-type strains with polyP of 700 units in length (p700). The YihI-SPA fusion protein was detected using an anti-Flag antibody. In contrast, for ZipA detection we used a commercially available antibody that we first validated in **Figure S1B.** In this assay YihI-SPA, but not ZipA, demonstrated the characteristic polyP shift indicative of polyP binding (**Figure 1C and 1D**) and this effect was dependent on the concentration of polyP used (**Figure S1C**).

### Characterization of the YihI PASK-like motif

Next, we aimed to further investigate how the YihI PASK was contributing to polyP binding. Previous work showed that mutation of lysine residues to arginine (K-R) abolished the polyP shift on NuPAGE gels for other PASK-containing proteins^20, 36, 50, 51^. Therefore, we used GST-YihI fusion proteins to test if this held true for YihI. Our bioinformatics analysis located the PASK motif to the C-terminus of the YihI protein (**Figure 2A**). Surprisingly, mutation of the two lysine residues in this region failed to prevent polyP interaction, suggesting that YihI does not bind to polyP via its defined PASK motif (**Figure 2B**). We noticed that the N-terminus of YihI is also lysine-rich (**Figure 2A**). Mutation of 7 N-terminal lysines to arginine severely abrogated the polyP shift (**Figure 2B**). Therefore, we conclude that YihI interacts with polyP primarily through this region. A YihI mutant where all lysines were replaced with arginine residues completely lost its ability to bind polyP as judged by NuPAGE analysis (**Figure 2B**), suggesting that other lysines may also contribute to polyP binding, at least in the absence of those in the N-terminus. While the N-terminus does not fit the formal definition of a PASK motif, it does contain a number of serine (3) and acidic (5) residues in addition to 7 lysine residues required for polyP interaction. Therefore, we refer to this region as ‘PASK-like’. Notably, both the N- and C-termini of YihI are disordered (**Figure 2C**), which is fitting for the molecular chaperone and scaffold-like functions of polyP^4, 52^. Additionally, the N-terminus of YihI may play regulatory functions. For example, truncation mutants lacking residues 1-45 show enhanced binding to Der and activation of Der’s GTP hydrolysis activity^48^, suggesting an overall inhibitory role for the N-terminus of YihI. We speculate that polyP binding may regulate these functions *in vivo*.

**Figure 2:**
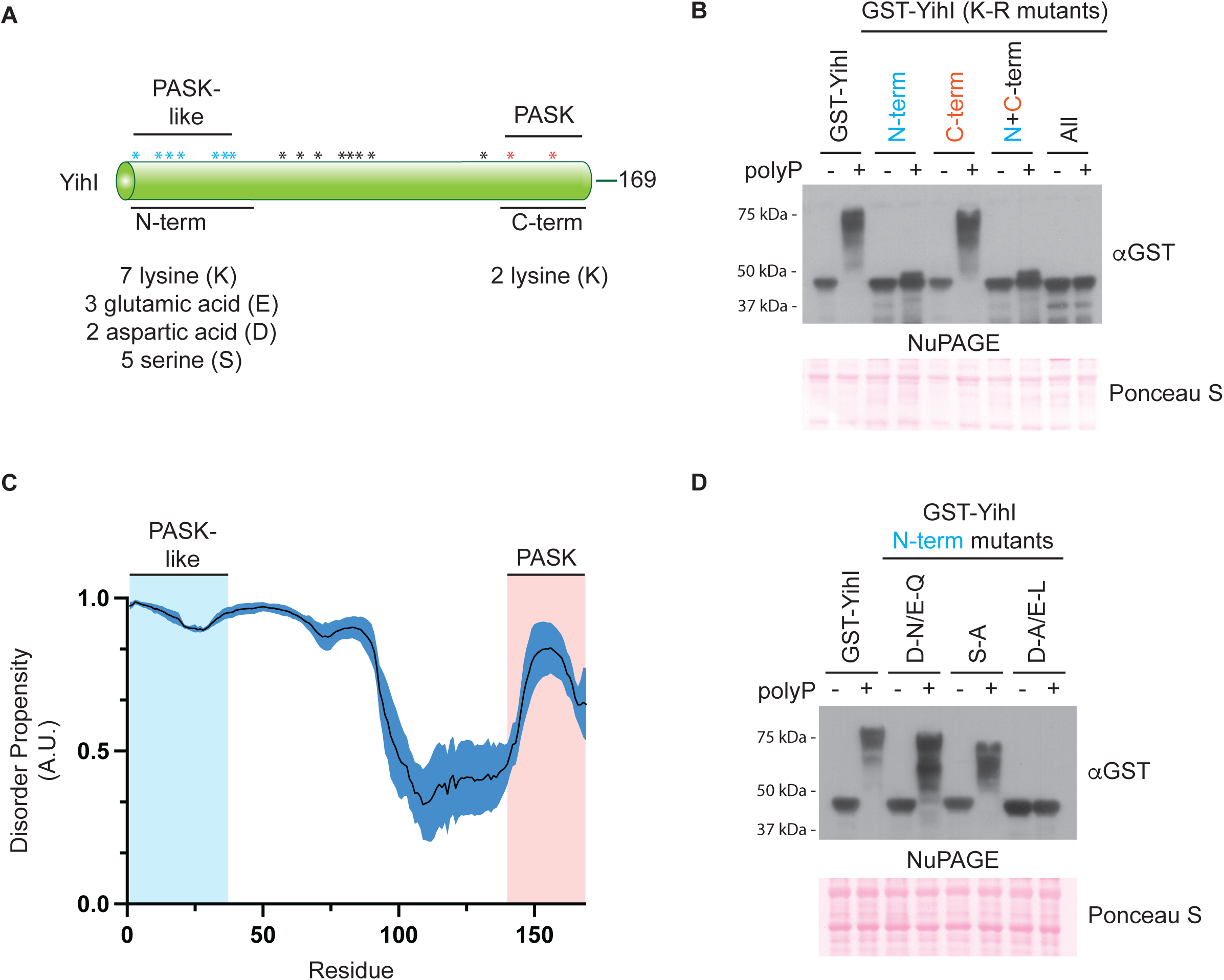
PolyP binds a disordered lysine-rich region of YihI. **(A)** Schematic and amino acid distribution of full-length YihI (169 residues total). YihI has a C-terminal PASK domain and a N-terminal PASK-like domain. The indicated amino acids distributed across the PASK or PASK-like domains were targeted for mutagenesis experiments. An asterisk (*) is used to display the distribution of PASK (orange), PASK-like (blue) and other lysine (black) residues within YihI. **(B)** PolyP binds primarily via the N-terminus of YihI. *In vitro* polyP binding assay was conducted (as described in Figure 1B) using whole cell extract expressing wild-type or lysine to arginine (K-R) mutated GST-YihI. **(C)** Disorder propensity of YihI shows that the N- and C-termini are highly unstructured (>0.5). Graph shows the average (±standard error) of computational prediction scores, represented as arbitrary units (A.U.), that were obtained using NetSurfP-3.0^95^, Metapredict^96^ and IUPred3^97^. **(D)** The N-terminal PASK amino acids play a structural role in promoting polyP binding. Various GST-YihI mutants were grown and analyzed as described in (B). D-N/E-Q = aspartic acid to asparagine/glutamic acid to glutamine; S-A = serine to alanine; D-A/E-L = aspartic acid to alanine/glutamic acid to leucine. For both (B) and (D), samples were resolved using NuPAGE, transferred to PVDF and probed using an anti-GST antibody. Ponceau S was used to show that samples migrated equally. Images are representative of results from ≥3 experiments.

Previous mutagenesis work on yeast targets demonstrated that serine residues in PASK motifs are not required for polyP binding^36^. In contrast, mutation of acidic residues (aspartic and glutamic acid) to alanine or leucine prevented polyP interaction^53^. In both cases, analogous N-terminal mutations resulted in a similar impact on polyP binding to GST-YihI (**Figure 2D**). To test if polyP binding depends on the negative charge of these acidic residues, we also mutated aspartic and glutamic acids to asparagine and glutamine, respectively. With these changes, GST-YihI was still able bind to polyP (**Figure 2D**), suggesting that negative charge *per se* is not required for polyP interaction, at least for this ‘PASK-like’ region of YihI (**See Discussion**).

### Novel non-PASK polyP-binding proteins in *E. coli*

To extend our search for polyP binding proteins in bacteria, we took advantage of two sets of *E. coli* strains where individual open-reading frames are expressed as fusion proteins with C-terminal SPA (781 strains) or TAP (243 strains) epitope tags (**Supplemental Table 1**)^54^. We generated protein extracts from these strains and carried out *in vitro* polyP binding assays, as described above (**Figure 1B**). The SPA tag^55^ contains a 3Flag epitope and the TAP tag has a protein A moiety that is recognized by most mouse antibodies. Therefore, we used a mouse anti-Flag antibody to detect both SPA and TAP targets after NuPAGE gel electrophoresis and western blotting.

After accounting for redundancy between the two epitope-tagged sets and proteins that were not detected by western blotting, we evaluated polyP binding for a total of 589 unique proteins using this assay (**Figure 3A** and **Supplemental Table 1**). Seven of these (1.2% of total proteins screened) shifted on NuPAGE gels in the presence of polyP (**Figure 3B**). With 4288 predicted open reading frames in *E. coli*, we anticipate at least ∼50 proteins from *E. coli* would undergo a polyP shift in this assay. This value is likely an underestimate, as work with human polyP interactors demonstrated that not all undergo polyP shifts on NuPAGE gels^32^. Indeed, we observed that the Lon protease, a well-characterized polyP binding protein from *E. coli*^38, 39^, does not shift on NuPAGE gels even in the presence of high concentrations of polyP (**Figure S2**).

**Figure 3:**
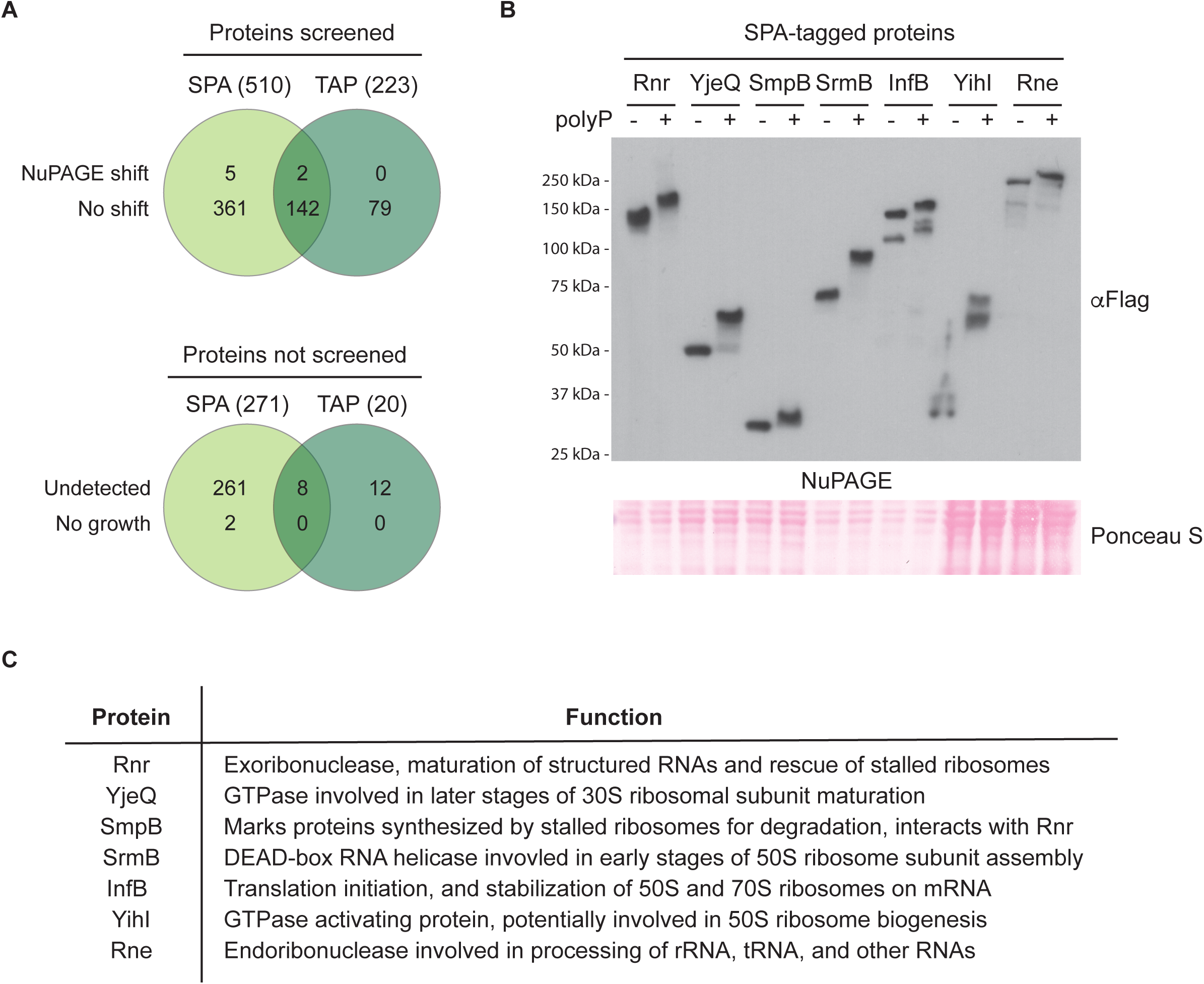
A screen for novel polyP-binding proteins in *E. coli.* **(A)** A total of 589 unique *E. coli* proteins were screened for polyP binding. Together, the SPA and TAP collection sets contain a total of 1024 strains with epitope tags encoded at the chromosomal loci of relevant open reading frames. Of these, 152 proteins are redundantly tagged between the SPA and TAP collection sets, and 291 (283 non-redundant) could not be screened for polyP binding. **(B)** Seven novel polyP binding proteins were identified by the screen. Proteins that shifted from the screen were reconfirmed using the *in vitro* polyP binding assay. Samples were resolved using NuPAGE, transferred to PVDF and probed using an anti-Flag antibody which detects the SPA tag. Ponceau S was used to show that samples migrated equally. Images are representative of results from ≥3 experiments. **(C)** The 7 polyP binding proteins are involved in ribosome biogenesis or translation processes. General descriptions of each protein’s functions are provided.

Intriguingly, all 7 polyP binders identified have links to ribosome assembly or function (**Figure 3C**). This finding is consistent with the enrichment of this same category in our yeast polyP-PASK interaction study^20^, as well as the remodelling of nucleoli, the site of ribosome biogenesis, in human cells ectopically expressing bacterial PPK to produce high levels of polyP^56^. Altogether, this suggests the possibility of evolutionarily conserved roles for polyP in the regulation of translation.

### Interaction of target proteins with endogenous polyP

To test if native polyP was also able to bind our newly identified targets, we switched SPA-tagged strains grown in LB media to MOPS minimal media to induce nutrient starvation and polyP accumulation prior to protein extraction and NuPAGE analysis. Out of the 7 targets, SrmB-SPA, and YihI-SPA consistently displayed an obvious MOPS-induced polyP shift while Rnr-SPA did so occasionally (**Figure S3A**). This result is perhaps surprising considering that the chain lengths of polyP that accumulate during MOPS appear to be larger than the p700 chains used in our *in vitro* assays (**Figure S3B**). We speculate that long-chain bacterial polyP is organized *in vivo* in a manner that in some instances hinders its interaction with protein targets. In support of this idea, we found that a large fraction of polyP that accumulates during MOPS treatment is resistant to ectopically expressed yeast Ppx1 (*Sc*Ppx1), a highly active exopolyphosphatase (**Figure S3C**). This finding is reminiscent of the situation in mammalian cell culture where *Sc*Ppx1 treatment results in a partial, but not complete, loss of the polyP signal in nuclear polyP foci detected using the PPBD-Xpress tag probe^57^. In contrast, *Sc*Ppx1 overexpression in yeast appears to completely degrade the non-vacuolar pool of polyP synthesized by *E. coli* PPK expression^58^.

### Functional interaction between *rnr* and *ppk*

We reasoned that some genes encoding polyP-interacting proteins might display genetic interactions with Δ*ppk* under conditions where polyP is important for cell growth or viability (**Figure S4A**). As previously reported and consistent with work from other groups^3, 59^, we found that Δ*ppk* mutants displayed a slow growth phenotype on MOPS minimal media relative to wild-type controls (**Figure 4A and Figure S4A**). This phenotype is likely attributable to an extended lag phase and decreased doubling time in Δ*ppk* mutant cells^42^. We observed that deletion of *rnr* does not impact polyP levels in wild-type cells (**Figure S4B**) but consistently improved the growth of Δ*ppk* mutant cells on MOPS media (**Figure 4A & S4A**). While the rescue was not complete, we conclude that in Δ*ppk* mutants, one or more activities of Rnr hinder cell growth during nutrient limitation.

**Figure 4:**
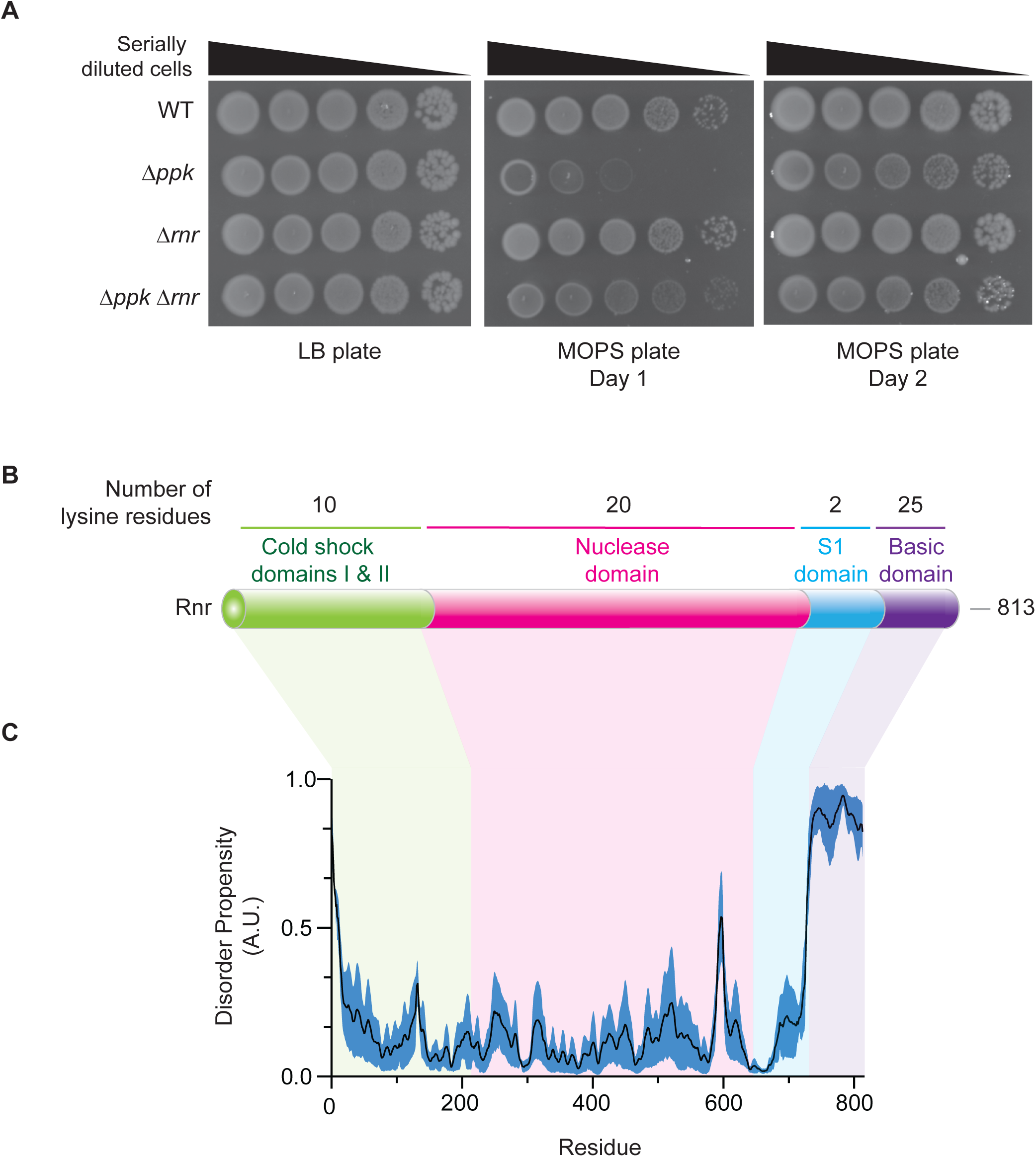
Rnr is functionally regulated by polyP. **(A)** Loss of *rnr* partially rescues the slow growth phenotype of *ppk* mutants. The indicated strains were serially diluted and spotted on LB or MOPS plates and incubated at 37°C as indicated. Images are representative of results from ≥3 experiments. **(B)** Schematic of the functional domains of full-length Rnr (813 residues total). Rnr has 2 cold shock domains (residues 1-216), a nuclease domain (residues 217-643), an S1 domain (residues 644-730) and a basic domain (residues 731-813). **(C)** The basic domain of Rnr has a high disorder propensity (>0.5). Graph shows the average (±standard error) of computational prediction scores, represented as arbitrary units (A.U.), that were obtained using NetSurfP-3.0^95^, Metapredict^96^ and IUPred3^97^.

Rnr (also referred to as VacB in literature) is a 3’ to 5’ exoribonuclease that plays a role in maintaining RNA homeostasis in cells^60, 61^. It primarily targets rRNAs and structured RNAs, including RNA duplexes, but not DNA^60, 62–64^. Rnr has a complex role *in vivo*. It is thought to play a role in RNA turnover and the recycling of excess rRNA during stress, such as starvation, cold shock, and stationary phase growth^65–67^. It has also been proposed to participate in trans-translation through its role in the maturation of tmRNA^68^, which binds SmpB (another polyP binding protein identified by our screen)^69^ and is required for tagging abnormal peptides and releasing stalled ribosomes^70–72^. Additionally, in an SmpB-dependent manner^73^, Rnr degrades ‘non-stop’ transcripts that result in ribosome stalling^72, 74^.

These complex functions and interactions of Rnr are mediated by various domains that work together in a coordinated manner. For example, Rnr possesses two cold shock domains with helicase activity^75^, cold shock specific functions^75^, and a role in substrate binding^62^, as well as a catalytic core termed the ribonuclease domain where reduction reactions take place^62, 76^ (**Figure 4B**). It also possesses S1 and basic domains that are involved in protein stabilization^77^, substrate positioning^62^, and ribosome binding^78^ (**Figure 4B**). Intriguingly, the basic domain is both disordered (**Figure 4C**) and lysine-rich, hinting at a possible role in polyP binding.

### Complex regulation of Rnr by PPK and polyP

To map the region of Rnr required for interaction with polyP, we expressed its individual domains as GST-fusion proteins and carried out *in vitro* polyP binding assays as described for YihI. These experiments demonstrated that the C-terminus of the protein (S1+basic domain) was responsible for polyP binding (**Figure S5A**). Indeed, deletion of this region from chromosomally expressed Rnr (detected using an anti-Rnr antibody, validated in **Figure S5B**), resulted in a loss of the polyP shift (**Figure 5A**), as did mutation of 27 S1+basic lysine residues to arginine (K-R) (**Figure 5B**). Since NuPAGE assays determine protein-polyP interactions under largely denaturing conditions, we also tested if polyP interacts with Rnr in its folded state. To do this, Rnr-3Flag was immunoprecipitated under non-denaturing conditions and incubated with polyP prior to washing and elution with sample buffer. In this experiment, unbound polyP is expected to be removed prior to NuPAGE analysis (**Figure S5C**). Immunoprecipitated Rnr incubated with polyP shifted on NuPAGE gels after washing, suggesting that polyP can also bind to Rnr when folded (**Figure S5C**).

**Figure 5:**
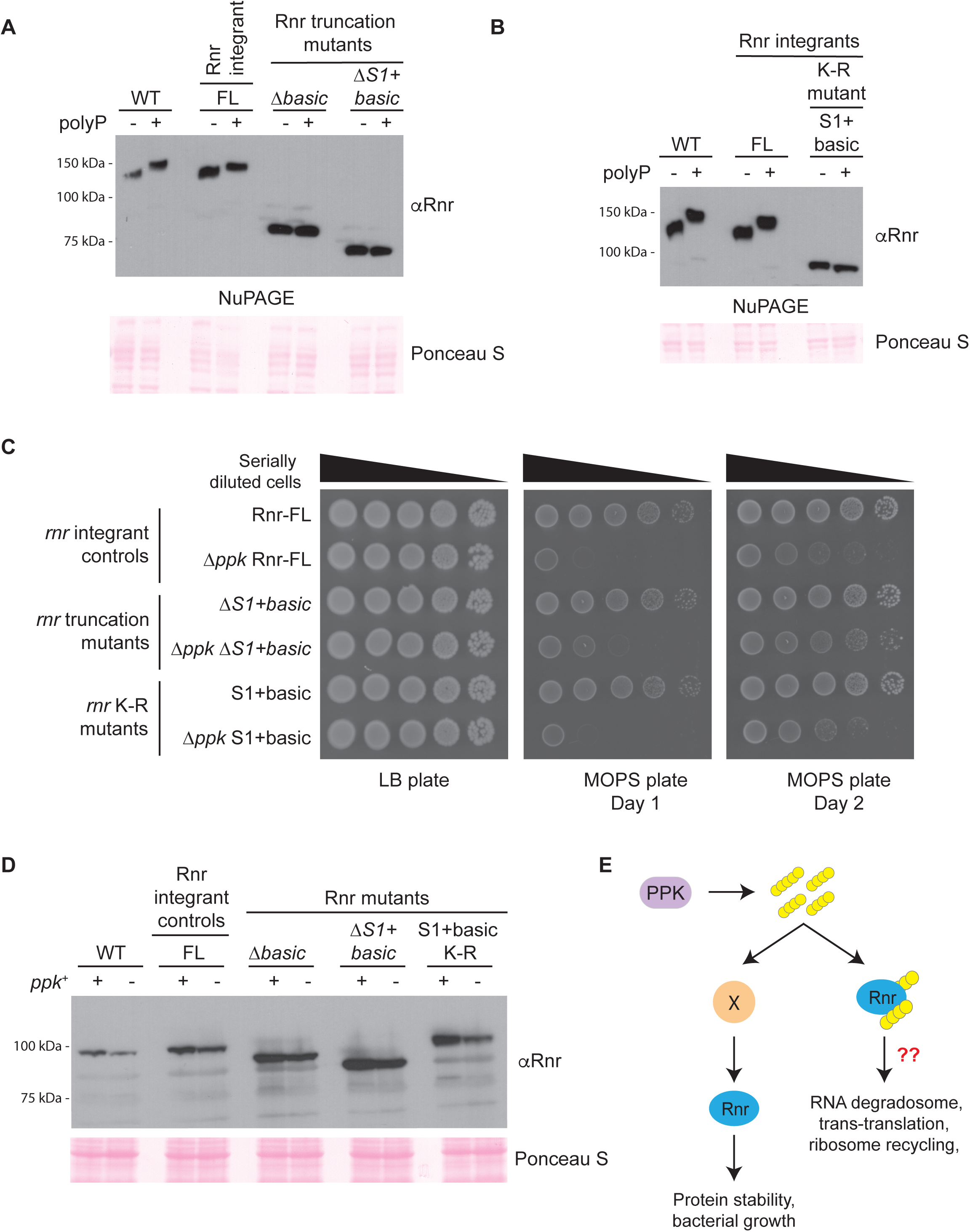
The Rnr S1 and basic domains are involved in polyP binding. **(A-B)** The characteristic NuPAGE shift is abrogated when the S1 and basic domains of Rnr are (A) truncated or (B) mutated. An *in vitro* polyP binding assay was conducted using the indicated chromosomally truncated or lysine to arginine (K-R) mutated Rnr strains. FL represents the wild-type Rnr protein expressed in a background that is isogenic to the truncated and mutated strains (see methods *Bacterial strains* section for details on how these strains were constructed). Samples were resolved using NuPAGE, transferred to PVDF and probed using an anti-Rnr antibody. Ponceau S was used to show that samples migrated equally. Images are representative of results from ≥3 experiments. **(C)** Truncation but not K-R mutation of the S1+basic polyP-binding domain partially rescues the slow growth phenotype of *ppk* mutants. The indicated strains were serially diluted and spotted on LB or MOPS plates and incubated at 37°C as indicated. Images are representative of results from ≥3 experiments. **(D)** Expression of wild-type and mutant Rnr is downregulated in *ppk* mutants compared to wild-type cells during growth in MOPS. FL is as described for (A). Whole cell extract from wild-type and mutant Rnr strains that were grown in LB media then exposed to nutrient down shift for 3 hours were resolved using 10% SDS-PAGE, transferred to PVDF and probed using an anti-Rnr antibody. Ponceau S was used to show equal loading. Images are representative of results from ≥3 experiments. **(E)** Model of how polyP impacts Rnr function. PolyP synthesized by PPK can indirectly impact Rnr stability through an unknown pathway. PolyP may also bind Rnr directly to modulate a variety of functions related to translation control and mRNA metabolism.

In growth assays, deletion of the Rnr S1+basic domain, but not the basic domain on its own, improved growth of Δ*ppk* mutants on MOPS (**Figure 5C** and **Figure S5D**), suggesting that together, these regions mediate toxicity in MOPS media in the absence of polyP. If polyP binding to the S1+basic domain functions to promote growth of wild-type cells on MOPS media, we predict that under these conditions, wild-type cells expressing the K-R mutant should display a slow growth phenotype, similar to Δ*ppk* mutants. However, the K-R mutant grew similarly to wild-type cells on MOPS media (**Figure 5C**).

Next, we investigated if polyP-binding could impact Rnr expression. In exponential phase Rnr is rapidly degraded as a result of acetylation at lysine544 (K544; within the S1 domain)^79^. This degradation is thought to be mediated through an interaction with SmpB via the Rnr C-terminus that results in recruitment of HslUV and Lon proteases^77, 80^. In contrast, Rnr is stabilized during stationary phase and under stress conditions^81^, and its activity increases upon carbon, nitrogen and phosphorus starvation^66^. Therefore, since protein binding to polyP has been shown to modulate protein degradation^39^, we evaluated Rnr expression in wild-type and Δ*ppk* mutants during nutrient starvation in MOPS, where polyP levels in wild-type cells are normally high. We found that Rnr levels were reproducibly decreased in Δ*ppk* mutants (**Figure 5D**). However, like the genetic interactions described above, this effect was not directly mediated by polyP binding to Rnr, because truncation and K-R mutants that failed to interact with polyP (**Figure 5A** and **5B**) still showed decreased expression in the Δ*ppk* mutant background (**Figure 5D**). Therefore, we conclude that PPK plays an indirect role in Rnr expression while the role of Rnr’s polyP binding remains unclear (**Figure 5E**).

## DISCUSSION

In this work, we have identified 7 novel polyP binding proteins in *E. coli* and provided additional evidence to support an evolutionarily conserved role for polyP in the regulation of protein translation.

Previous work in eukaryotic cells has demonstrated that PASK motifs, often found in predicted disordered regions, are a frequent site for polyP binding. To our surprise, PASK-containing proteins are rare in bacterial models commonly used for polyP research. Of the 7 proteins that we identified as polyP binders, only YihI has a PASK motif that fits our previous definitions of 75% D/E/S/K with at least one lysine in a 20 amino acid window. However, this motif makes at most a minor contribution to YihI’s polyP binding activity. Instead, we mapped this function to the N-terminus of the protein which we refer to as PASK-like. Similar to what was described for PASK motifs, mutation of YihI’s N-terminal lysine residues to arginines abolished polyP interaction as judged by the NuPAGE shift assay, whereas mutation of serines to alanines had no effect. Since arginine holds a greater positive charge than lysine, these experiments demonstrate that concentration of positive charge is not the sole determinant of polyP binding. On the other hand, mutation of the acidic residues, glutamic and aspartic acid to leucine or alanine, abolished polyP interaction. However, mutation of these same residues to uncharged glutamine and asparagine had no effect. Our interpretation of these data is that the negative charge of the PASK-like region is dispensable for polyP interaction. One possibility is that the glutamic and aspartic acid residues instead provide a structural context for polyP binding with surrounding lysine residues and that this unique context is preserved with the glutamine and asparagine substitutions. We surmise that the impact of these substitutions will also hold true for canonical PASK motifs in proteins such as yeast Nsr1 and Top1^36^, but this remains to be tested. Indeed, it is currently unclear if there is a tangible difference between PASK and PASK-like motifs in the way that they interact with polyP molecules. In addition to PASK motifs, polyP has also been shown to bind other linear motifs namely polyHistidine^35^ and polyLysine stretches^34^. However, these motifs do not appear to be present in the targets that we identified. Certainly, we cannot discount the possibility that other linear polyP interacting motifs exist, and it will therefore be important to systematically map the binding region for each target.

A limitation of our work is that our detection of polyP-protein interactions relied on the previously described NuPAGE ‘polyP shift’ assay. Not all polyP binding proteins shift upon NuPAGE analysis in the presence of polyP, as demonstrated with our experiments using purified Lon protease. As such, we have no doubt that additional polyP-binding proteins in *E. coli* remain to be identified. In particular, polyP interactions that require folded protein structures would be missed in our assay, as these would likely be denatured during NuPAGE analysis. On the other hand, the ability of targets identified here to interact with polyP under denaturing conditions suggests polyP may play a role in their folding or re-folding after cellular stress. Indeed, this chaperone-like activity for polyP has been described previously for the *E. coli* CsgA protein involved in biofilm production^28^, and globally to stabilize proteins that become insoluble in response to oxidative stress^4^.

Our work adds to a growing body of evidence for an evolutionarily conserved role for polyP in regulating protein translation. Previously, Δ*ppk* mutant strains were found to have disrupted polysome profiles^82^. Further, polyP promotes translation fidelity *in vitro* and Δ*ppk* mutants have increased mistranslation *in vivo*^82^. In this regard, it is noteworthy that all of our newly identified polyP-binding proteins are linked in some way to ribosome biogenesis or translational control. We speculate that polyP binding to these proteins may therefore play a role in reprogramming translation during stress. Alternatively, polyP may help to stabilize critical regulators of translation so that they are ready to act upon a return to favourable growth conditions. Most of our new targets are highly conserved across other bacterial species (**Supplemental Table 2**), and in some cases are expressed in pathogens^83–87^. Therefore, it will be important to test whether polyP interacts with their homologs in these species.

In addition to identifying novel polyP-binding proteins linked to translation, we found evidence for a bidirectional regulation between PPK and Rnr. Namely, while PPK promotes Rnr expression during starvation, Rnr is also detrimental for growth in Δ*ppk* mutant cells grown on MOPS media. We propose a model where Rnr’s various molecular functions must be carefully balanced during cellular stress, and that this balance is lost in the absence of PPK. Importantly, mutation of the S1+basic domain lysine residues of Rnr to prevent polyP binding did not impact Rnr protein levels or growth characteristics in an otherwise wild-type background. The simplest explanation for these observations is that the described bidirectional regulation is indirect in that it does not depend on the Rnr-polyP interaction. Alternatively, the regulation may become relevant under situations where growth is already compromised, as is the case in Δ*ppk* mutant cells.

Since polyP binds to Rnr in both its denatured and folded states, it is possible that polyP impacts Rnr biology at multiple levels, and additional work will be required to tease out specific molecular functions. For example, based on the C-terminal binding of polyP, it may disrupt functions associated with the S1 and basic domains, which includes an intrinsically disordered segment. As observed for Nsr1 and Top1 in yeast^36^, polyP binding may disrupt the interaction between tmRNA-SmpB and Rnr, which is mediated via the basic domain^77^. This in turn could play a role in stabilizing Rnr in some contexts, potentially through cross regulation with a previously reported acetylation at K544 that is known to promote Rnr turnover^79^. Additionally, polyP interaction with the S1 domain could alter Rnr substrate selectivity^64^. Very likely, polyP binding to Rrn is part of a broader function for polyP in adapting to cellular stress. We note, for example, that Rnr also plays a role in the RNA degradosome in conjunction with Rne^88^, another polyP-binding protein identified in our screen. As such, we do not discount the possibility that dramatic phenotypes would only be observed after mutating polyP binding motifs on multiple proteins involved in the processes of ribosome biogenesis or translation.

*In vivo*, local subcellular distribution of polyP may govern whether a protein interacts with it. Moreover, we demonstrate here that a large fraction of intracellular polyP is resistant to degradation via overexpression of the highly active yeast Ppx1. As such, *in vivo* some polyP may be inaccessible to potential protein interactors. It is tempting to speculate that this property is dictated by the ability of polyP to phase separate *in vivo* and the investigation of this property and its relationship to protein-polyP interactions is deserving of further attention. Another important area for future investigation will be to determine how polyP-protein interactions are reversed upon return to normal growth conditions. We speculate that bacterial PPX enzymes may play a critical role in this process. Indeed, this activity has been demonstrated previously for yeast Ppx1^36^. Alternatively, in the presence of ADP, PPK itself may drive the conversion of protein-bound polyP to ATP. This would relieve polyP dependent modulation of translation, while providing ATP pools required for renewed efforts towards ribosome biogenesis and growth.

## MATERIALS & METHODS

### General information about strains and plasmids

All bacterial strains and plasmids, as well as their sources, used in this work are listed in **Supplemental Table 3**. Plasmids were sequenced using Sanger sequencing (Genome Quebec) or Nanopore sequencing (Plasmidsaurus). All plasmids (**Supplemental Table 3**) generated for this work will be made available from Addgene (www.addgene.com) upon final publication. The sequences of oligonucleotides used for cloning or genetic manipulations are available upon request.

### Bacterial strains

Unless otherwise indicated, the MG1655 strain background was used for all experiments. All lab-generated strains used in this study are listed in **Supplemental Table 3**. The Dharmacon Collection of SPA- and TAP-tagged *E. coli* strains (DY330 background) were obtained from Horizon Discovery and have been described previously^54^. The strains list for both of these collection sets can be found online under the *Resources* tab (https://horizondiscovery.com/en/non-mammalian-research-tools/products/e-coli-tagged-orfs#description).

Chromosomally-tagged and deletion strains were generated using the lambda-red homologous recombination system using pKD46 (induced with 0.2% arabinose)^89^ or pSIM6 (induced with a temperature shift to 42 °C for 12 minutes)^90^. Respectively, the kanamycin deletion and 3Flag-kanamycin tagging cassettes were amplified from pKD4^89^ and pSUB11^91^ plasmids. Rnr truncations were made by using forward primers that introduced a premature stop codon and led to recombination that deleted the end of the gene, replacing the region with the KanR selection cassette. For the basic and S1+basic mutant, a stop codon was introduced after residue 2190 and 1929, respectively. An FRT scar was also introduced at the end of full-length Rnr to control for polar effects^92^. This strain is referred to as full-length (FL) and is isogenic to the *rnr* truncation mutants (ΔS1+basic and Δbasic). For genetic experiments, cells were made electrocompetent and plasmids or double-stranded DNA used for recombineering were transformed into cells via electroporation^93^. Antibiotics were added when appropriate: kanamycin (50 μg/ml) and ampicillin (100 μg/ml). As needed, the resistance markers used for selection of positive transformants were removed using the pCP20 FLP-recombinase system^94^. Epitope tag insertions and deletions were confirmed by PCR, followed by western blotting.

### Plasmids

The GST-YihI wild-type and mutant sequence plasmids were purchased from GenScript. The respective YihI sequences were cloned between the EcoRI and NotI sites. These vectors are called: pYihI, pYihI-N-term K-R, pYihI-C-term K-R, pYihI-N+C-term K-R, pYihI-All K-R, pYihI-N-term D-N/E-Q, pYihI-N-term S-A, pYihI-N-term D-A/E-L.

The Rnr cold shock domains I and II (residues 1-216), nuclease domain (residues 217-643) and S1+basic domains (residues 644-813) were cloned into pGEX4T1 between the EcoRI and NotI restriction sites using Gibson Assembly Cloning. These vectors are called pRnr-CSD, pRnr-ND and pRnr-S1BD, respectively.

The *ScPPX1* plasmid was constructed by amplifying the S. cerevisiae *PPX1* sequence from pET-15b-His-*PPX1* and cloning it between EcoRI and SalI sites of pBAD18. The empty and cloned vectors are called pBAD18 and p*ScPPX1*, respectively.

### Bacterial growth conditions

#### General growth conditions

SPA- and TAP-tagged strains^54^ (DY330 background) were grown at 30°C while all other strains in the MG1655 background were grown at 37°C. Unless otherwise specified, strains were grown in LB media.

#### Nutrient downshift

Starvation experiments were performed as previously described^42^. Briefly, overnight cultures grown in LB were diluted to 0.1 OD_600_ in LB media and grown to mid-exponential phase (∼0.6 OD_600_) before being switched to MOPS minimal media. Cells were pelleted and washed once with 1x PBS to remove trace LB before resuspension in freshly prepared MOPS media (1x MOPS – Teknova, 0.1 mM K_2_HPO_4_, 0.4% glucose). Cells were grown in MOPS media for the indicated amount of time. Typically, we see peak polyP accumulation after 3 hours in MOPS media. For western blotting and polyP extractions, 3 and 5 OD_600_ equivalent of cells were harvest by centrifugation, respectively.

### PASKMotifFinder Software

The PASKMotifFinder software used to search for PASK motifs was implemented in Java and is platform independent. The code is open source and freely available at the following GitHub repository: https://github.com/LavalleeAdamLab/PASKMotifFinder/. The software uses a sliding window approach to scan subsequences of 20 amino acids throughout the proteome and identifies regions where D, E, S and/or K amino acids make up at least 75% of the window (i.e. 15 amino acids), and contain at least one K. The program was run on the *E. coli* (strain K12) proteome UP000000625, and other proteomes listed in **Supplemental Data 1**. All proteomes were downloaded on January 13, 2025 from UniProt release version 2024_06.

### *In vitro* polyP binding assay

*In vitro* assays were conducted using whole cell extract or purified proteins.

Whole cell extract was prepared as described under *Western blotting – protein extraction* using 200 µL of overnight culture. For the polyP binding assay, 10 µL of whole cell extract was incubated at room temperature in the presence of 10 mM sodium phosphate pH 6.0 (control matching the pH of the polyP) or p700 (Kerafast) polyP for 20 minutes. For the concentration shift assay, control reactions contained sodium phosphate matching the highest concentration of polyP used. All control (minus polyP) and reaction samples were boiled for 10 minutes and loaded onto a NuPAGE Bis-Tris Mini Protein Gel, 4-12%, 1.5 mm. See methods on *Western blotting* for the subsequent steps used for visualizing proteins.

Purified Rts1 and Lon protease (purchased from SinoBio) were used for the *in vitro* polyP binding assay, as described previously^20^. Briefly, the purified proteins (0.032 mg of each) were incubated at room temperature with increasing concentrations of p700 (5, 10, 15 and 20 mM) or 20 mM sodium phosphate pH 6.0 (negative control) for 20 minutes. All control (minus polyP) and reaction samples were boiled for 10 minutes and loaded onto a NuPAGE Bis-Tris Mini Protein Gel, 4-12%, 1.5 mm. The gel was stained using the Invitrogen Colloidal Blue Staining Kit.

### Western blotting

#### Protein extraction

As indicated, 200 µL of an overnight culture or 1.5-3 OD_600_ equivalents of cells were harvested by centrifugation for analysis by western blotting. Cells were resuspended in 100 µL of sample buffer (800 uL sample buffer stock (160 mM Tris-HCl pH 6.8, 30% glycerol, 6% SDS, 0.004% bromophenol blue) + 100 uL 1 M DTT, 100 1.5 M Tris-HCl pH 8.8), boiled for 10 minutes and then centrifuged at 13,000 rpm for 2 minutes to remove insoluble material. The supernatant was transferred to a fresh tube. Typically, to normalize for equal loading 10 µL and 13 µL of extract from wild-type and Δ*ppk* mutant cells were loaded per blot, respectively.

#### Gel electrophoresis and transfer

NuPAGE or SDS-PAGE gels were used to resolve protein extracts. SDS-PAGE was primarily used to visualize protein levels while NuPAGE gels were solely used to detect polyP dependent shifts of our candidate proteins. After electrophoretic separation, proteins were transferred onto PVDF membranes and visualized by western blotting using the indicated antibodies. SDS-PAGE and NuPAGE buffer recipes have been described previously^20^.

#### Western blotting

Membranes were blocked for 20 minutes with shaking using 5% milk in TBST and washed 3 times for 10 minutes after both primary and secondary antibody incubations. See **Supplemental Table 4** for incubation conditions for each antibody. Note: both SPA- and TAP-tags were detected using an anti-Flag antibody, which was then detected using a goat anti-mouse secondary coupled to HRP. After probing, target proteins were detected using Immobilon Western Chemiluminescent HRP Substrate and exposure to autoradiography film from Thomas Scientific. Scanned images were opened in Photoshop, and linear brightness and contrast adjustments were made to lighten the image background. Adjustments were applied evenly across the entire image prior to cropping and labelling. For all western blots, staining with Ponceau S was used to verify equal loading, protein migration, and even transfer across the PVDF membrane.

### Mapping polyP binding domains

YihI (wild-type and mutated sequences) and Rnr domains were cloned into pGEX4T1 and transformed into BL21 for expression. Overnight cultures harboring the plasmids were diluted 1/100 in LB + ampicillin and grown to mid-exponential phase (∼2 hours). Cells were induced with 0.1 mM IPTG for 2 hours and 1.5 OD_600_ equivalent of cells were harvested. Whole cell extract was prepared by resuspending pellets in 100 µL of sample buffer and was used to conduct *in vitro* polyP binding assays (described above).

### YihI and Rnr disorder predictions

The amino acid sequences for YihI (UniProt accession: B1XAM2) and Rnr (UniProt accession: P21499) were entered into the following disorder prediction programs: NetSurfP-3.0 (https://services.healthtech.dtu.dk/services/NetSurfP-3.0/)^95^, Metapredict online (v3.0) (https://metapredict.net/)^96^ and IUPred3 (https://iupred3.elte.hu/)^97^. These programs were selected to account for variations between prediction algorithms and prevent bias^98, 99^. Disorder scores from the three programs were averaged and graphed with the standard error envelope using GraphPad Prism as described by Pastic *et al*.^100^. See **Supplemental Data 2** for individual prediction scores.

### Screen for polyP binding proteins

#### Bacterial growth

SPA and TAP collection sets^54^ were pinned on LB + kanamycin plates and grown overnight at 30°C. The next day, grown colonies were inoculated into 3 mL LB + kanamycin and grown at 30°C overnight.

#### Protein extraction

From the overnight cultures, 200 µL of cells were pelleted and used to prepare whole cell extract as described in the *Western blotting* section of the methods.

#### *In vitro* polyP binding assay

The extracts were screened as described above. In brief, whole cell extract of the tagged strains was incubated in the absence or presence of polyP (modal size p700) and resolved using NuPAGE (as described in the *Western blotting* section of the methods).

#### Confirming positive hits

Positive candidates were streaked for single colonies and the correct position of the tag was confirmed via PCR analyses. Primers used in these confirmation assays are available upon request. These proteins were re-screened using sodium phosphate pH 6.0 (matching the pH of p700) as a control. With the exception of the screen, sodium phosphate pH 6.0 was used as a control for all *in vitro* polyP binding assays. The screened strains are listed in **Supplemental Table 1.**

### Ppx1 overexpression assay

Strains harboring the pBAD18 (empty vector) and *ScPPX1* plasmids were grown in the presence of ampicillin at all stages. Overnight cultures were diluted into LB media and induced with 0.5% arabinose, grown to mid-exponential phase and then nutrient downshifted into MOPS media (as described above under *Growth conditions* for 3 hours. The only variation is that for the MOPS media, 0.5% arabinose was included, and glucose (0.4%) was replaced with glycerol (0.5%) as the carbon source. For western blotting and polyP extraction, 3 and 5 OD_600_ equivalent of cells were harvested, respectively.

### PolyP extraction

#### Extraction

PolyP extractions were performed as described previously^42^ and have been briefly summarized with similar wording here. Five OD_600_ equivalents of cells were used for polyP extractions. Cell pellets were resuspended in LETS buffer (100 mM LiCl, 10 mM EDTA, 10 mM Tris-HCl pH 7.4 and 0.2% SDS). PolyP was extracted using the phenol/chloroform method and precipitated overnight at −20°C in 100% ethanol containing 120 mM sodium acetate. Precipitated polyP was pelleted by centrifugation, resuspended in 30 μL sterile water and stored at −80°C.

#### Gel analysis

Extracted polyP, mixed 1:1 with loading dye (10 mM Tris-HCl (pH 7), 1 mM EDTA, 30% glycerol, and bromophenol blue) was resolved using a 15.8% TBE-urea gel (5.25 g urea, 7.9 ml 30% acrylamide, 3 ml 5xTBE, 150 μl 10% APS, and 15 μl TEMED) run at 100 V for 1 hour and 45 mins in 1x TBE. The gel was then stained in fixing solution (25% methanol, 5% glycerol) containing 0.05% toluidine blue and then de-stained in fixing solution without toluidine blue. For the polyP standards, 6 µL of each chain length at the specified concentration, p130 (1.25 mM) and p700 (1 mM), was mixed 1:1 with loading dye and 10 µL was loaded into the gel.

### Growth assays

Spot tests were conducted as described previously, with the details reiterated here^42^. The indicated strains were streaked on LB plates and incubated overnight at 37°C. The next day, single colonies were resuspended in 100 µL of sterile water and serially diluted 10-fold 5 times in sterile water. Next, 5 µL of each dilution was spotted onto LB or MOPS (1x MOPS, 0.4% glucose, 0.1 mM K_2_HPO_4_) and incubated at 37°C. To prepare MOPS plates, 10x MOPS, glucose and K_2_HPO_4_ were added after autoclaving water + agar. LB plates were imaged after 1 overnight while MOPS plates were imaged post-day 1 and −day 2. Spot tests were imaged using ImageQuant LAS 4000 and edited across the entire image by making minor linear brightness and contrast adjustments in Photoshop to lighten the background.

### Immunoprecipitation and *in vitro* polyP-binding assay

Overnight culture of the Rnr-3Flag tagged strain was diluted to 0.1 OD_600_ in 300 mL of LB and grown to mid-exponential phase. Fifty OD_600_ equivalent of cells were harvested on ice and stored at −80°C until used for immunoprecipitation. Cells were resuspended in 700 µL buffer A (50 mM HEPES pH 7.9, 150 mM NaCl, 1 mM EDTA, 0.5% Triton X-100, 5% glycerol, 1 mM PMSF, Roche cOmplete protease inhibitor cocktail tablet) and sonicated (Misonix 3000) on ice for 3 cycles of 10 seconds at power level 3 with 30 second rest in between. The lysate was cleared by centrifugation for 15 minutes at 15,000 rpm at 4°C and then incubated with 5 µL of the 50% anti-FLAG M2 magnetic bead slurry (Sigma Aldrich M8823-1ML) for 1 hour at 4°C. Next, the beads and bound proteins were washed 3 times with 1 mL of buffer A using cut pipette tips. Beads were then resuspended in 10 mM sodium phosphate (pH 6.0) or p700 in a final volume of 250 uL of buffer B (same as buffer A, but with 0.05% Triton X-100 and no protease inhibitor tablet) and incubated at room temperature with end-to-end rotation for 20 minutes. Excess polyP was then washed away using three 1 mL washes with buffer B. Finally, proteins were eluted in 60 µL 2X sample buffer containing no DTT by incubating at 65°C for 10 minutes. Finally, the sample was transferred to a new tube, DTT was added to a final concentration of 100 mM and the sample was boiled at 100°C for 10 minutes prior to resolving (20 µL per sample) on NuPAGE gels.

### Chromosomal Rnr lysine to arginine mutants

Gene fragments encoding lysine to arginine mutants for the S1+basic domain were purchased from Twist Bioscience. Towards the 5’ and 3’ ends, the fragments had homology needed for the recombineering transformation and homology towards the beginning of the pKD4 cassette, respectively. In a separate PCR reaction, the KanR cassette was amplified from pKD4. This reaction used forward and reverse primers introducing homology towards the K-R fragments and homology needed for the recombineering transformation, respectively. Next, in a two-step PCR the two fragments (K-R gene fragments + KanR cassette) were combined at a 1:1 molar ratio and amplified. The final products were gel extracted and transformed into *rnr*-Δbasic mutants by electroporation. Correct integration of the K-R mutations was confirmed by Premium PCR sequencing from Plasmidsaurus.

### Anti-GST antibody purification

The anti-GST antibody was purified from sera collected from rabbits injected with a GST-Cdc26 fusion protein^101^. Prior to anti-Cdc26 antibody purification on a Cdc26 affinity column, the sera was cleared of anti-GST antibodies on a 50 mL GST affinity column, as described^101^. Anti-GST antibodies were eluted with 100 mM glycine, pH 2.1, neutralized in 2 M tris-base and dialyzed in antibody storage buffer (1x PBS, 500 mM NaCl, 50% glycerol).

### Anti-Rnr antibody

An antibody towards Rnr was raised by immunizing New Zealand NZW female rabbits with purified GST-Rnr nuclease domain (amino acids 649-1929). This domain was chosen for immunization as it does not bind polyP and therefore, would not impact detection of truncated or mutant Rnr. The pGEX4T1 vector was cloned with sequence encoding the Rnr nuclease domain (amino acids 649-1929) using standard Gibson assembly. The oligonucleotides used for this strategy are available upon request. The vector was transformed into BL21 DE3 pLysS *E. coli* and plated on LB + ampicillin + chloramphenicol. Overnight cultures of cells harboring the vector were diluted to 0.1 OD_600_ and grown at 30°C until they reached OD_600_ of 0.4-0.6. Cells were then induced with 0.25 mM IPTG for 4 hours prior to harvesting and freezing at −80°C in 40 mL of freezing buffer (1x PBS).

#### GST-fusion purification

Frozen cell lysates were thawed in a water bath, then immediately transferred onto ice to prevent degradation. Next, 40 mL of 1x PBS containing 2 mM EDTA, 2 mM EGTA, 2 mM PMSF, 30 mM DTT, 1 M NaCl was added to bring the final volume of the cell slurry to 80 mL. Lysozyme was added at a final concentration of 200 µg/mL and the lysate was incubated on ice for 30 minutes before disruption using the Misonix sonicator (3 cycles of 1 min at power level 7, with 2 minutes rest on ice in between). Triton X-100 was added after sonication to a final concentration of 0.5%. The lysate was centrifuged at 40,000 x g for 30 minutes, then cleared through a 0.45 micron filter and then batch bound for 2 hour to glutathione agarose beads that had been equilibrated in wash buffer with detergent (1x PBS, 0.1% NP-40, 0.5 M NaCl, 1 mM DTT, 1 mM EDTA, 1 mM EGTA and 1 mM PMSF). The GST-bound beads were washed with 15 column volumes of wash buffer with detergent and 5 column volumes with wash buffer without detergent (no NP-40). The column was eluted with 20 mL elution buffer (50 mM Tris-HCl pH 8.0, 0.5 M NaCl, 10 mM glutathione, 1 mM DTT and 1 mM PMSF) and dialyzed into the storage buffer (1x PBS, 100 mM NaCl, 15% glycerol). The dialysis buffer was changed 3 times. The GST-purified protein was concentrated using an Amicon Ultra-15 centrifugal filter and quantified against serially diluted BSA standards using gel electrophoresis and Coomassie staining and quantification. The final protein was aliquoted at stored at −80°C.

#### Injection preparation

For the first immunization, 675 µL of purified protein, at a final concentration of 2 mg/mL, was combined with 75 µL penicillin (10 U/mL)/streptomycin (10 µg/mL) and 750 µL Freund’s Complete Adjuvant. In contrast, Freund’s Incomplete Adjuvant was used to prepare subsequent booster injections.

#### Injections

For both primary and booster immunizations, four 100 µL subcutaneous and two 250 µL intramuscular injections were administered. The booster was administered 4 weeks after the initial immunization and the rabbits were terminally bled 4 weeks later. All antigen injections and blood collections were administered under general anesthesia to minimize pain and suffering. Rabbits were first sedated with injectable sedatives butorphanol and midazolam and induced into general anesthesia with inhaled isoflurane from a precision vaporizer.

#### Antibody validation

Sera was validated in the lab by western blotting, using wild-type and Δ*rnr* (negative control) *E. coli* whole cell extracts (**Figure S5B**).

### Sequence similarity comparison analyses

Protein sequences of the 7 hits were individually blasted against the *Salmonella enterica* (taxid: 28901), *Helicobacter pylori* (taxid: 210), *Streptomyces coelicolor* (taxid: 1902) and *Mycobacterium tuberculosis* (taxid: 1773) proteomes using NCBI Blast (https://blast.ncbi.nlm.nih.gov/Blast.cgi?PAGE=Proteins). Search settings were as follow: standard databases, non-redundant protein sequences (nr) and blastp (protein-protein BLAST). For each comparison, a single result with the greatest query coverage and, secondly, lowest E-value is presented in **Supplemental Table 2**.

## ACKNOWLEDGEMENTS

This work was funded by a Canadian Institutes of Health Research (CIHR) Project Grant to MD and MLA (PJT-148722). KB was supported in part by an Ontario Graduate Scholarship. IA was supported in part by a Canada Graduate Scholarship from the Natural Sciences and Engineering Research Council of Canada (NSERC). We thank members of the Downey lab for helpful suggestions and critical reading of the manuscript.

## ETHICS STATEMENTS

This study was performed in strict accordance with standards for animal care and use outlined in the Canadian Council on Animal Care (CCAC) policies and guidelines. The University of Ottawa holds a certificate of Good Animal Practice with the CCAC and is a registered research facility under the Province of Ontario’s Animals for Research Act. The animal use protocol (BMIe-3469) was approved by the University of Ottawa Animal Care Committee.

---

**Supplemental Figure 1:**
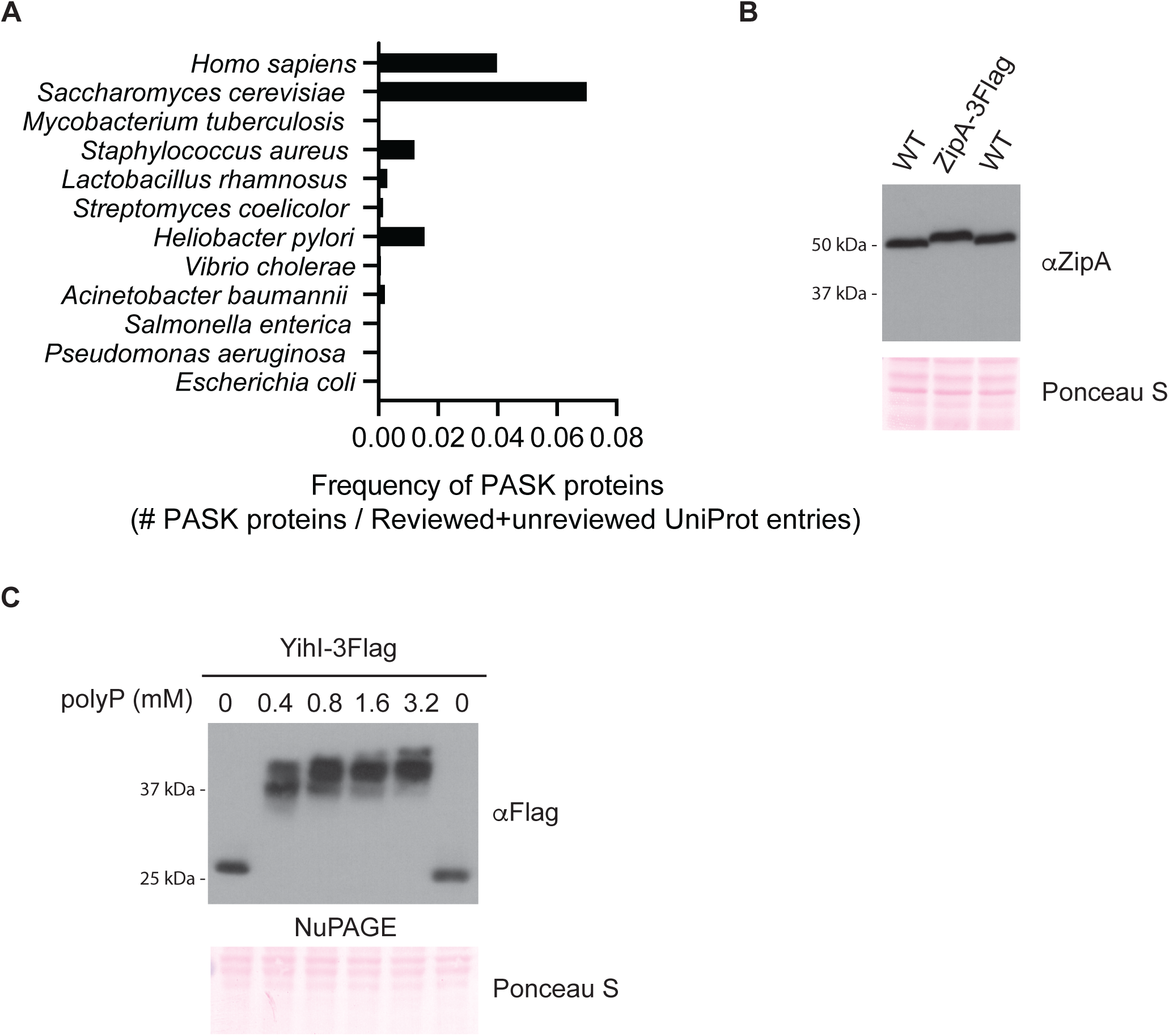
The PASK is not a good indicator of polyP-protein binding in bacteria. **(A)** PASK frequency using reviewed+unreviewed proteomes. The number of proteins containing 1 or more PASK motifs (75% D/E/S/K content with at least one lysine within a 20 amino acid window) from reviewed and unreviewed proteomes of the indicated species were normalized by the total number of reviewed+unreviewed UniProt entries for each species. **(B)** Anti-ZipA antibody validation blot. Whole cell extracts from wild-type and ZipA-3Flag tagged strains were resolved on 12% SDS-PAGE, transferred to PVDF and probed using an anti-ZipA antibody. Ponceau S was used to show equal protein loading and that samples migrated equally. A deletion mutation of Δ*zipA* could not be used because *zipA* is essential. ZipA-3Flag displayed a shift in migration as expected, confirming the antibody recognizes ZipA. **(C)** YihI-polyP binding shifts on NuPAGE are polyP concentration dependent. Whole cell extract from a YihI-3Flag tagged strain was used for an *in vitro* polyP binding assay in the presence of increasing concentrations of polyP. Samples were resolved using NuPAGE, transferred to PVDF and probed using an anti-Flag antibody. Ponceau S was used to show that samples migrated equally. Images are representative of results from ≥3 experiments.

**Supplemental Figure 2:**
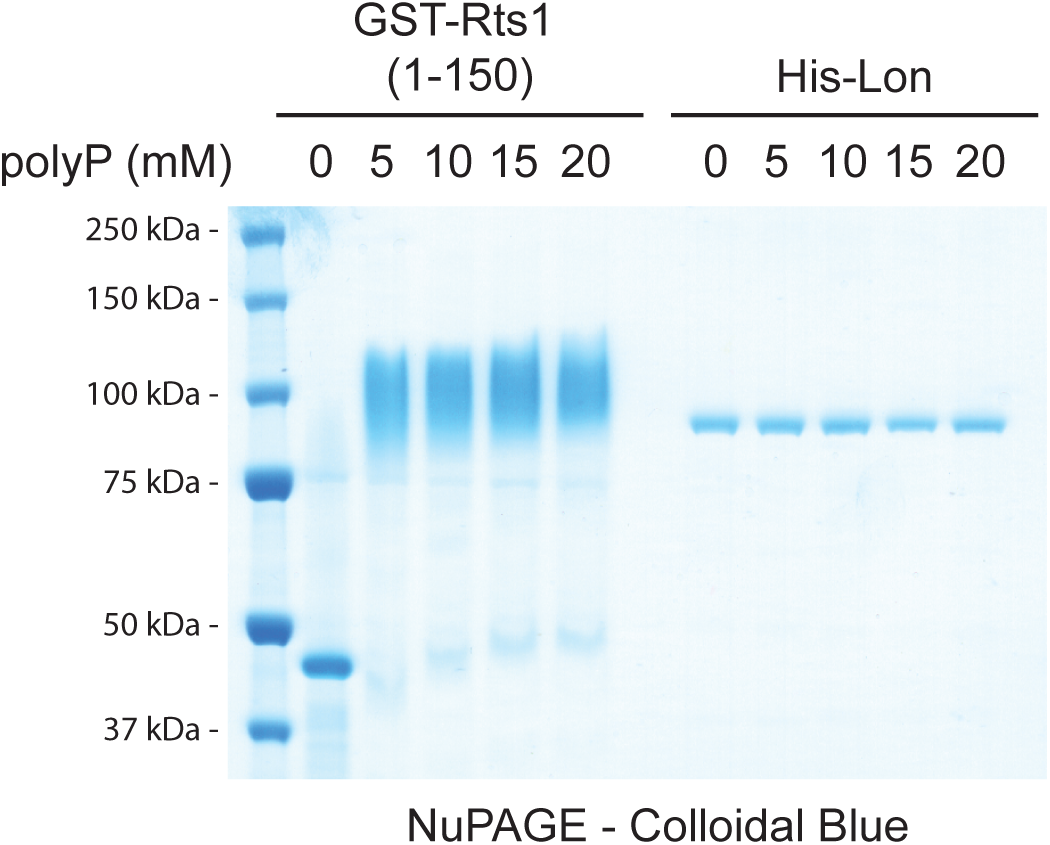
NuPAGE electrophoresis is not effective for detecting polyP binding to Lon. Lon does not display the characteristic polyP binding shift on NuPAGE gels. Increasing concentrations of polyP (p700) were incubated with 0.032 mg of purified Rts1 (positive control)^20^ or Lon protease. Samples were resolved using NuPAGE and the gel was stained using Colloidal Blue to visualize the proteins. Image is representative of results from ≥3 experiments.

**Supplemental Figure 3:**
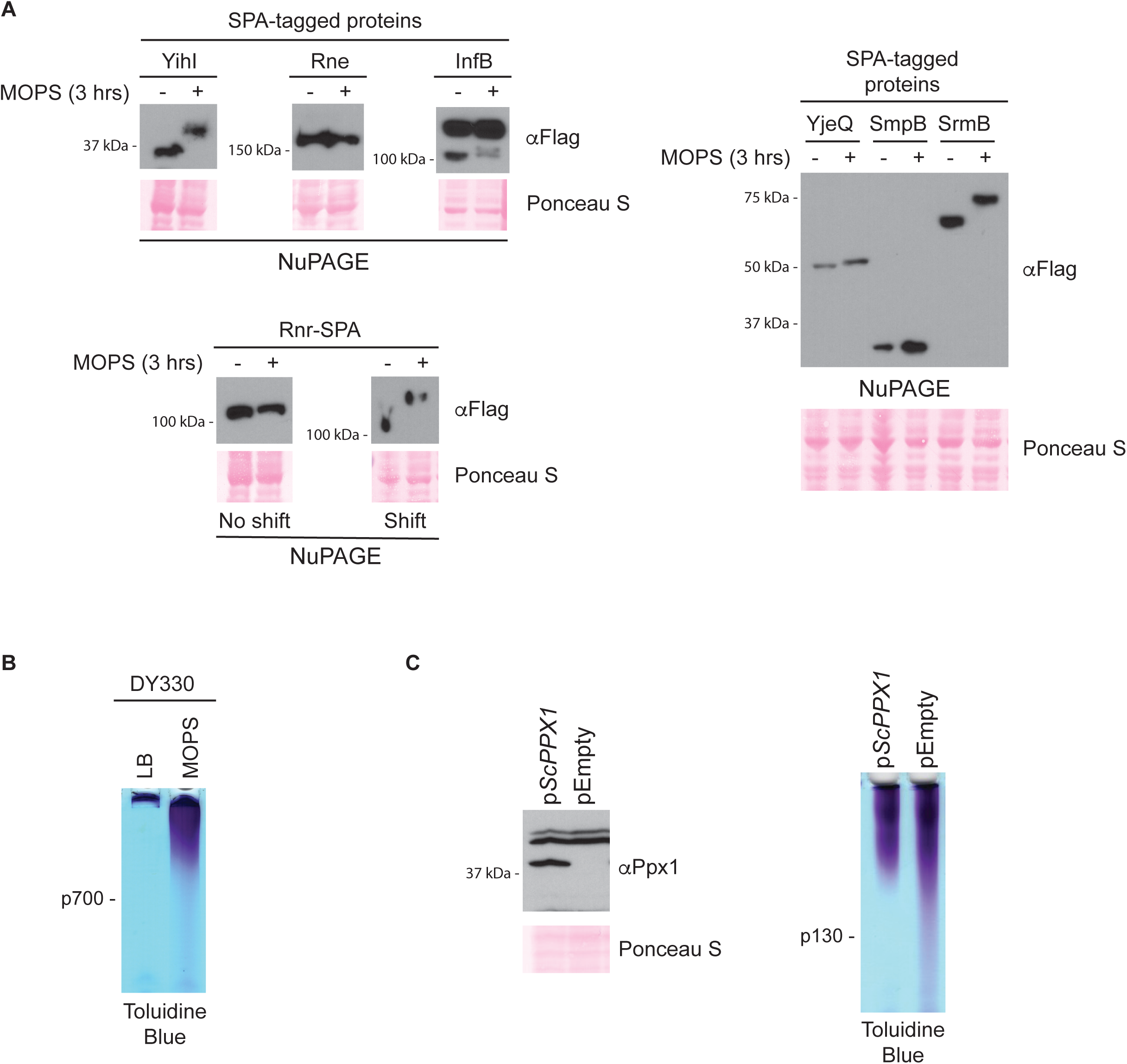
PolyP binding proteins may have limited access to endogenous polyP that accumulates in response to stress. **(A)** YihI, SmpB and Rnr display NuPAGE shifts in the presence of endogenous polyP. Whole cell extract from SPA-tagged strains that were grown in LB media (-MOPS) or exposed to nutrient downshift (+ MOPS) for 3 hours were resolved using NuPAGE, transferred to PVDF and probed using an anti-Flag antibody which detects the SPA tag. Ponceau S was used to show that samples migrated equally. Images are representative of results from ≥3 experiments. **(B)** *E. coli* makes long chain polyP after 3 hours in MOPS media. PolyP was extracted from cells grown in LB media or MOPS for 3 hours (as described for S3A) was resolved using a TBE-urea acrylamide gel and stained using toluidine blue. The gel shows that endogenous polyP is longer than the p700 standard. Images are representative of results from ≥3 experiments. **(C)** Endogenous polyP is not fully degraded by ectopic expression of *S. cerevisiae* exopolyphosphatase Ppx1 (*Sc*Ppx1). Western blotting (left) and polyP extractions (right) of *E. coli* with p*ScPPX1* or the empty vector. Cells were grown in LB media in the presence of 0.5% arabinose (the inducer) before undergoing a nutrient downshift to MOPS media for 3 hours. PolyP and whole cell extracts were resolved using a TBE-urea acrylamide gel or 12% SDS-PAGE, respectively. The polyP gel was stained using toluidine blue and Ppx1 expression was detected using an anti-Ppx1 antibody. Images are representative of ≥3 results.

**Supplemental Figure 4:**
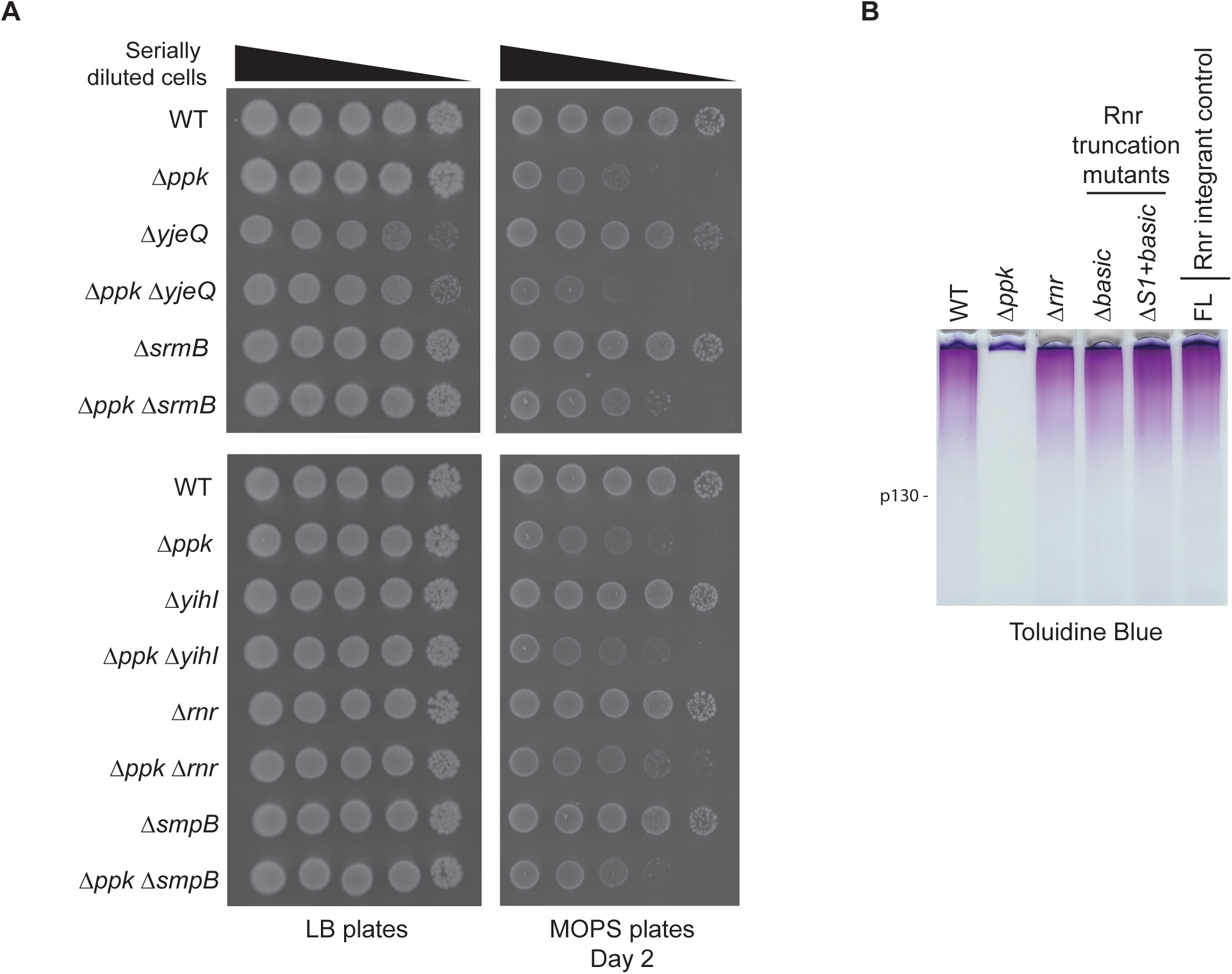
Loss of polyP binding proteins SrmB and Rnr rescues *ppk* mutant growth phenotypes. **(A)** Spot test of *E. coli* mutated for genes encoding the polyP binding proteins. The indicated strains were serially diluted and spotted on LB or MOPS plates and incubated at 37°C as indicated. Images are representative of results from ≥3 experiments. **(B)** Mutation of *rnr* does not impact polyP accumulation in an otherwise wild-type background. PolyP extracted from cells grown in LB media and exposed to nutrient down shift for 3 hours was resolved using a TBE-urea acrylamide gel and stained using toluidine blue. The migration of a standard of modal length p130 is indicated. FL represents the wild-type Rnr protein expressed in a background that is isogenic to the truncated and mutated strains (see methods *Bacterial strains* section for details on how these strains were made). Images are representative of results from ≥3 experiments.

**Supplemental Figure 5:**
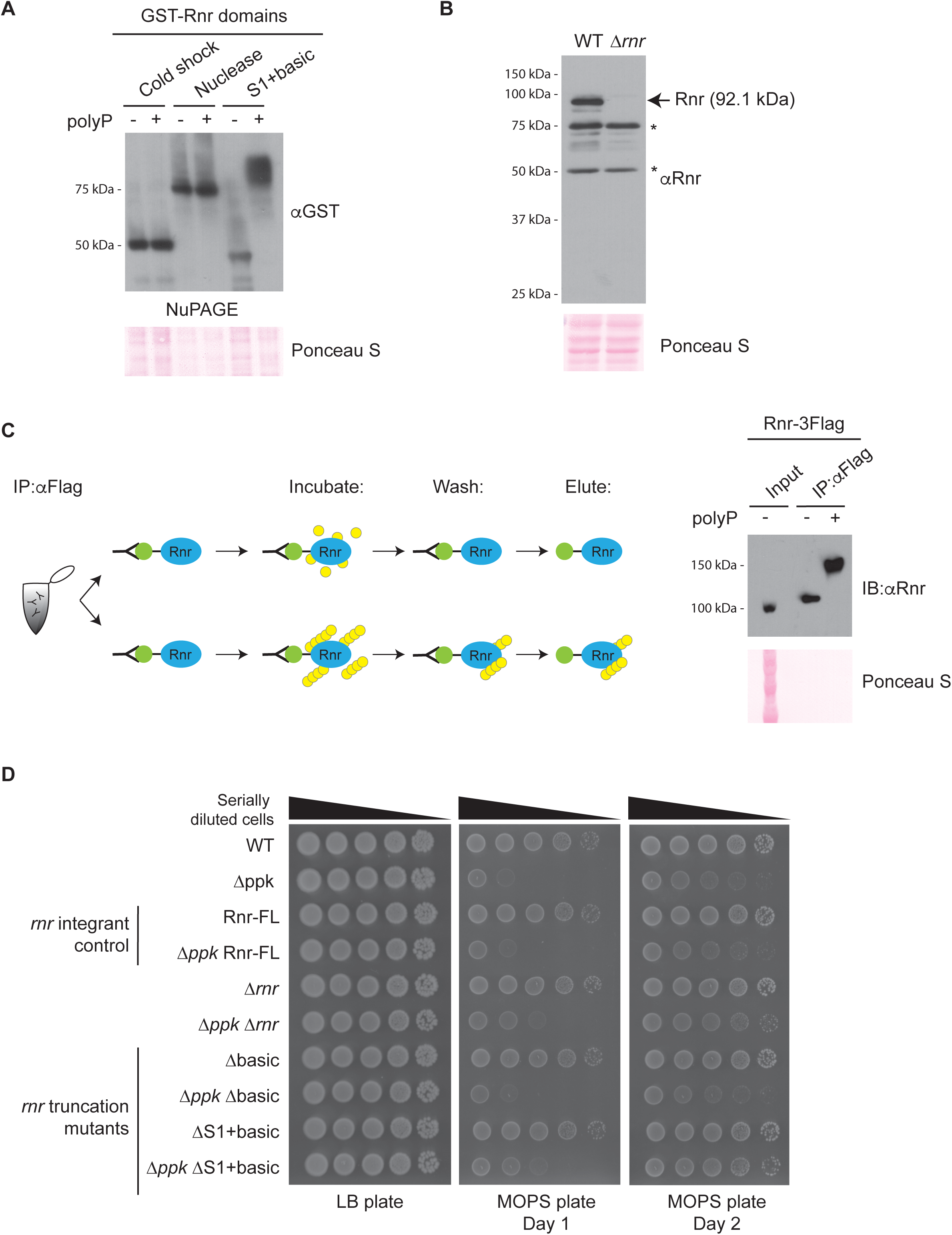
A complex interplay between PPK and the Rnr polyP binding domain. **(A)** PolyP binds to the S1 and basic domain of Rnr. Whole cell extract from cells expressing GST-tagged Rnr domains were used to conduct *in vitro* polyP binding assays, resolved using NuPAGE, transferred to PVDF and probed using an anti-GST antibody. Ponceau S was used to show that samples migrated equally. Images are representative of results from ≥3 experiments. **(B)** Anti-Rnr antibody validation blot. An arrow is used to show the band corresponding to Rnr while asterisks (*) indicate background bands. Whole cell extract from WT and Δ*rnr* strains was resolved on 12% SDS-PAGE, transferred to PVDF and probed using the anti-Rnr antibody. Ponceau S was used to show equal protein loading. **(C)** PolyP binds the native form of Rnr. Schematic (left): Rnr-3Flag was immunoprecipitated (IP) from whole cell extract under non-denaturing conditions using anti-Flag beads and then incubated with polyP (p700). Excess polyP was then washed away before eluting the protein. IP’ed proteins were resolved using NuPAGE, transferred to PVDF and probed using an anti-Rnr antibody. Images are representative of results from ≥3 experiments. **(D)** Loss of the Rnr S1 and basic domains rescues *ppk* mutant growth phenotypes comparable to Δ*ppk* Δ*rnr* double mutants. FL represents the wild-type Rnr protein expressed in a background that is isogenic to the truncated and mutated strains (see methods *Bacterial strains* section for details on how these strains were made). The indicated strains were serially diluted and spotted on LB or MOPS plates and incubated at 37°C as indicated. Images are representative of results from ≥3 experiments.

**Supplemental Table 1 (attached)**

Screened and unscreened strains from the SPA- and TAP-collection sets.

**Supplemental Table 2 (attached)**

Conservation analysis of polyP-binding proteins across species commonly used for polyP research.

**Supplemental Table 3 (attached)**

Bacterial strains and plasmids used in this study.

**Supplemental Table 4 (attached)**

Antibodies used in this study.

**Supplemental Data 1 (attached)**

Raw data used to determine frequence of proteins with a PASK motif in bacteria in (Figures 1A and S1A)

**Supplemental Data 2 (attached)**

Raw data used to graph YihI (Figure 2C) and Rnr (Figure 4C) disorder propensity graphs.

